# Cross-transmission of resistant gastrointestinal nematodes between wildlife and transhumant sheep

**DOI:** 10.1101/2023.07.21.550073

**Authors:** Camille Beaumelle, Carole Toïgo, Rodolphe Papet, Slimania Benabed, Mathieu Beurier, Léa Bordes, Anaïs Brignone, Nadine Curt-Grand-Gaudin, Mathieu Garel, Justine Ginot, Philippe Jacquiet, Christian Miquel, Marie-Thérèse Poirel, Anna Serafino, Eric Vannard, Gilles Bourgoin, Glenn Yannic

## Abstract

Wild and domestic ungulates can be infected with the same species of gastrointestinal parasitic nematodes. These parasites have free-living stages in the environment that contribute to the ease of transmission among different host species. In addition, gastrointestinal nematodes have developed resistance to anthelmintics which is now considered a major problem for the livestock sector. In a context where wild and domestic ungulates share the same pastures, the maintenance and circulation of resistant gastrointestinal nematodes between species have rarely been explored.

In the European Alps, domestic sheep are driven to high-altitude summer pastures and live in sympatry with wild ungulates for several months each year. In this study, we investigated the nemabiome of domestic sheep and Alpine ibex, *Capra ibex*, in three different areas of the French Alps to evaluate parasite circulation between the two host species. The Alpine ibex is a protected mountain ungulate that is phylogenetically related to sheep and hosts nematode species common to sheep.

Using internal transcribed spacer 2 (ITS-2) nemabiome metabarcoding, we found sheep and ibex share similar gastrointestinal nematodes, except for a few species such as *Marshallagia marshalli* and *Trichostrongylus axei*. This suggests that the long-term co-occurrence of sheep and ibex on mountain pastures has promoted the exchange of gastrointestinal nematodes between the two hosts. Based on the sequencing of the isotype 1 of the beta tubulin gene, associated with benzimidazole resistance, we found resistant nematodes in all sheep flocks and in all ibex populations. Our results demonstrated that ibex can host and shed resistant strains before transhumant sheep arrive on pastures, and thus could act as a refuge or even contribute to maintaining resistant gastrointestinal nematodes. The relative role of ibex in the maintenance and circulation of resistant strains in sheep remain to be determined.

## Introduction

Parasites represent a large proportion of animal diversity and are key components of food webs (Hudson et al., 2006). They are also essential determinants of the health, fitness, population dynamics and community composition of their hosts (Tompkins et al., 2011). Parasites of the Nematoda phylum infect a wide range of species worldwide, including animals and plants (Blaxter and Koutsovoulos, 2015). In animals, gastrointestinal nematode parasites are of major concern for livestock productivity and security as they can impact animal health and reduce animal production, with significant economic losses (Charlier et al., 2020; Roeber et al., 2013). Ungulates are usually infected by free-living larvae of gastrointestinal nematodes when they graze pasture. Infective larvae may survive several months in the environment depending on species and climatic conditions (O’Connor et al., 2006). Following ingestion, larvae complete their development to reach the adult stage in the digestive tract. Egg-laying occurs 2─4 weeks post-infection. The duration of infection by nematodes varies depending on species but lasts for at least 2 months from L3 ingestion (Deplazes et al., 2016). The larvae can arrest their development in the host (hypobiosis) during harsh climatic conditions, delaying egg-laying (Deplazes et al., 2016).

To limit parasite load and its impact on livestock health, the use of anthelmintics to treat livestock against gastrointestinal nematodes is a common and cost-effective practice (Vercruysse et al., 2018). Nonetheless, the repeated use of anthelmintics has led to the selection of anthelmintic-resistant strains of gastrointestinal nematodes. Resistance to several families of anthelmintics (e.g., benzimidazole, macrocyclic lactones, and levamisole) has been observed, and multi-resistance is increasing (Bordes et al., 2020; Kaplan and Vidyashankar, 2012; Rose et al., 2015; Rose Vineer et al., 2020).

In particular, resistance to benzimidazoles is widespread throughout the world (Kaplan and Vidyashankar, 2012), and is particularly common on sheep farms in Europe (Papadopoulos et al., 2012; Rose Vineer et al., 2020). Contrary to other anthelmintic families, the mechanisms of resistance to benzimidazoles are well known and documented (Whittaker et al., 2017); in resistant nematodes, specific mutations of the β-tubulin isotype-1 gene have been correlated with resistance (Charlier et al., 2022). Furthermore, large-scale screening based on molecular tools is now feasible for this resistance (Avramenko et al., 2019), whereas the recommended method in livestock (i.e., the fecal egg count reduction test; Kaplan et al., 2023) for the diagnosis of resistance to other anthelmintics requires techniques that are difficult to achieve in wildlife living in remote locations.

Some generalist gastrointestinal nematodes can infect several host species (Walker and Morgan, 2014), including both domestic and wild ungulates (e.g., Beaumelle et al., 2022; Cerutti et al., 2010). The transmission of gastrointestinal nematodes among hosts, even if they do not simultaneously occupy the same pastures, is possible thanks to their free-living infective larval stage, which may remain active in the environment for several months (Carlsson et al., 2013; Fiel et al., 2012; Walker and Morgan, 2014). Transmitted parasites can also include gastrointestinal nematodes resistant to anthelmintics. For instance, benzimidazole-resistant nematodes have been detected in free-living populations of roe deer (*Capreolus capreolus*), living in sympatry with livestock (Chintoan-Uta et al., 2014; Nagy et al., 2017). To date, the role of wild ungulates in the epidemiology of resistant nematodes remains to be determined, but it has been suggested that wildlife may act as a reservoir of resistant nematodes for livestock (Brown et al., 2022; Chintoan-Uta et al., 2014; Francis and Šlapeta, 2023; Laca Megyesi et al., 2019; Walker and Morgan, 2014). To accurately evaluate this hypothesis, we need to investigate the presence of resistant nematodes in co-grazing wild and domestic ungulates in different contexts (i.e., different host species, different landscapes, and under different climatic conditions).

Transhumant pastoralism is a common practice in the European Alps and consists in the seasonal movement of grazing livestock from lowland areas to mountain meadows in summer which provide fresh pasture for domestic ungulates, i.e., mainly sheep, but also cows or goats (Biber, 2010). Mountainous areas are inhabited year-round by wild ungulates, particularly those living at high altitude in the European Alps, like Alpine ibex (*Capra ibex*), or Northern chamois (*Rupicapra rupicapra*). While wild ungulates tend to spatially or temporarily avoid domestic herds during the summer (Acevedo et al., 2008), certain factors may contribute to their use of the same pastures.

Spatial segregation between wild and domestic ungulates is usually observed once livestock arrive on pasture (Brivio et al., 2022; Ryser-Degiorgis et al., 2002). Livestock are generally released onto the best grazing areas during the summer season (Chirichella et al., 2014; Richomme et al., 2006; Ryser-Degiorgis et al., 2002). Mountain ungulates have been observed to preferentially use the same grazing areas both before and after livestock use (Brivio et al., 2022; Ryser-Degiorgis et al., 2002). The presence of wild and domestic ungulates in attracting zones such as salt licks, even if not simultaneous, offers ample opportunities for parasite transmission. Therefore, these areas are considered hotspots for parasite infection (Richomme et al., 2006; Ryser-Degiorgis et al., 2002; Utaaker et al., 2023).

Consequently, transhumant pastoralism represents a risk for pathogen transmission between wild and domestic ungulates in mountain areas (Rossi et al., 2019). Pathogen exchange at the interface of wild and domestic ungulates have already been well documented. The Alpine ibex has been identified as the wildlife reservoir of brucellosis (*Brucella melitensis*) which was transmitted to cattle and humans in the Bargy massif in northern French Alps (Marchand et al., 2017). In addition, sheep have been confirmed as the domestic reservoir of Border disease, which caused a major viral outbreak in Southern chamois (*Rupicapra pyrenaica*) populations in the Pyrenees (Luzzago et al., 2016). The transmission of gastrointestinal nematodes has already been described between wild ungulates and transhumant domestic ungulates in mountainous areas (Cerutti et al., 2010; Citterio et al., 2006; Khanyari et al., 2022; Zaffaroni et al., 2000). However, no study has yet investigated the transmission of anthelmintic-resistant nematodes in a transhumant pastoral system.

In this study, we investigated the community of gastrointestinal nematodes infecting Alpine ibex and domestic sheep (*Ovis aries*) and the prevalence of resistance to benzimidazole in three different regions of the French Alps. The Alpine ibex was close to extinction at the beginning of the 19^th^ century but the reinforcement of its populations by several reintroductions in different parts of the Alps has increased the species’ overall abundance and range (Brambilla et al., 2022). Today, the Alpine ibex population is estimated at 52 000 individuals in Europe (Brambilla et al., 2020).

Ibex usually host species-specific gastrointestinal nematodes (Walker and Morgan, 2014) but they may also be exposed to generalist nematodes deposited by other related ungulate (i.e., Northern chamois, *Rupicapra rupicapra*) or domestic species (e.g., sheep). Furthermore, anthelmintic treatments are frequently applied to livestock by farmers to reduce parasite load and consequently the diversity of nematodes in sheep. Therefore, we expected significant differentiation in the nemabiome between the two species across the three studied mountain areas, with higher nematode diversity in ibex compared to sheep (H1). We also expected sheep to host benzimidazole-resistant strains of gastrointestinal nematodes, consistent with the general pattern observed for sheep in France (Papadopoulos et al., 2012; Rose Vineer et al., 2020). Thanks to the reintroduction programs in the second half of the 20^th^ century, ibex colonized pastures traditionally grazed by sheep. Consequently, we expected that ibex would also host benzimidazole-resistant gastrointestinal nematodes, albeit to a lesser extent, as resistance does not confer a selective advantage for nematodes in the ibex environment (Hahnel et al., 2018) (H2). Given the rarity of documented ibex dispersal events among the three studied mountain areas (Brambilla, 2020 R. Papet, C. Toïgo and E. Vannard, personal communication), we predicted genetic differences in nematode species/communities or strains (ASV: Amplicon sequence variant) among ibex populations due to genetic drift (H3).

## Materials and Methods

### Study areas

Samples of sheep and ibex feces were collected in the French Alps in three different mountain areas (Figure 1). The Belledonne mountain is located in the western part of the Alps in southeast France. The Cerces and Champsaur mountains are situated in the northern and the southern parts of the Ecrins National Park, respectively (Figure 1).

**Figure 1:**
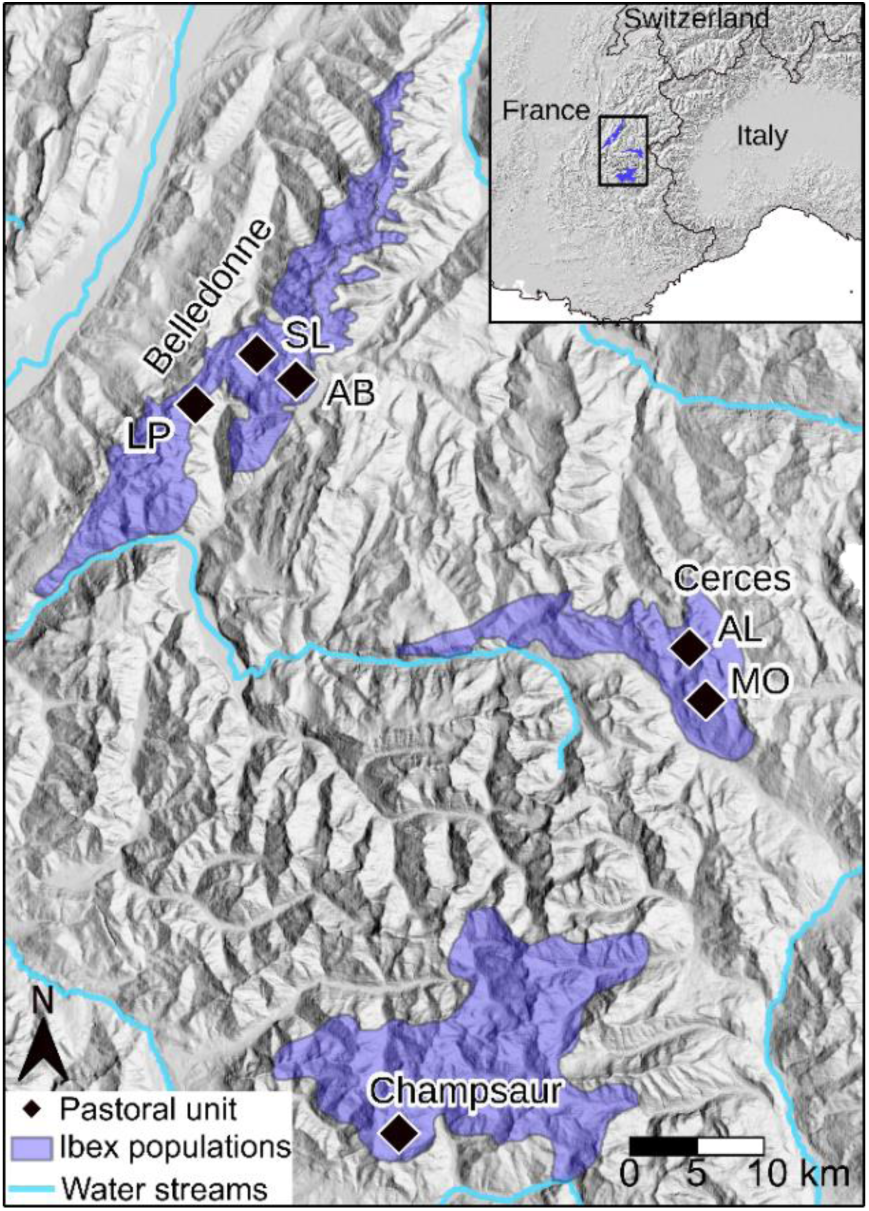
Map of the study area illustrating the three mountain areas (Belledonne, Cerces, Champsaur) in the French Alps where fecal samples from sheep and ibex were collected. In the Belledonne and Cerces areas, several pastoral units were sampled: Belledonne: AB: Ane Buyant, LP: La Pesée and SL: Sept Laux, Cerces: AL: Aiguillette de Lauzet, and MO: Montagne de l’Oule.

The three mountain areas are characterized by the presence of steep slopes, high peaks (>2500m) and agropastoral activities. Climatic conditions are harsh in these mountains with a mean temperature in winter (December-March) 2015-2019 of 0°C in Belledonne (alt:1785m), 1°C in Champsaur (alt:1620m) and 1.5°C in Cerces (alt:1553m). During summer (June-September), the mean temperature is 13°C in Belledonne, 14°C in Champsaur and 16°C in the Cerces (Réseau d’Observation Météo du Massif Alpin; www.romma.fr). Champsaur, the southern study area, has a Mediterranean influence. Consequently, rainfall is less important in this area compared to Cerces and Belledonne. Across areas, the vegetation is distributed along an elevation gradient from coniferous woodland (*Abies alba* and *Picea abies* in Belledonne and *Larix decidua* and *Pinus sylvestris* in Cerces and Champsaur) in the lower range of ibex, to a landscape dominated by heathland with *Rhododendron ferrugineum*, *Vaccinium* spp. and *Juniperus communis*, and grassland (*Carex* spp. *Festuca* spp.) above the tree line (Ozenda, 1985).

The ibex populations were established in Belledonne in 1983 with the introduction of 20 ibex, in Cerces, in 1959-1961 with the introduction of 6 ibex, and in Champsaur in 1994-1995 with the introduction of 30 ibex. Traditional pastoral activity is practiced in all massifs, where sheep flocks arrive from the plain on foot or by truck in early summer to graze mountain pastures.

In the Belledonne mountain, the distribution of ibex range between 630m and 2860m over 200km². Population size is estimated at 800 individuals. The size of the sheep herds are 750 ewes followed by their lambs in La Pesée, 900 ewes in Sept Laux, and 1600 ewes in Ane Buyant. A dozen rams are also present within the La Pesée and Sept Laux herds, as well as some goats in La Pesée.

The ibex population located in the Cerces mountain is estimated at 320 individuals and occupies an area of 120km², between 1410m and 3100m. In Cerces, the sheep herd located in the West (Aiguillette du Lauzet) includes 3 breeding farms for a total of 800 sheep and the herd located in the East (Montagne de l’Oule) includes 4 breeding farms for a total of 940 sheep (Table 1).

**Table 1:**
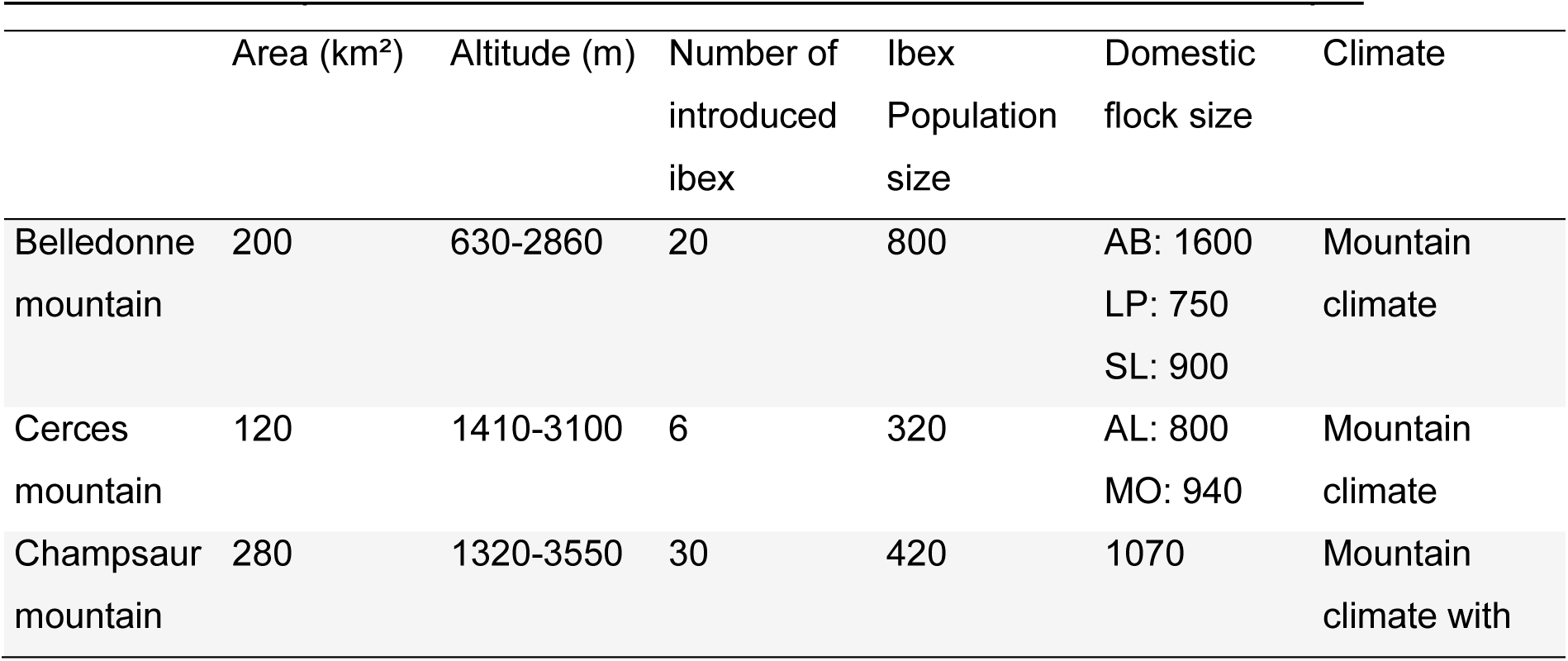
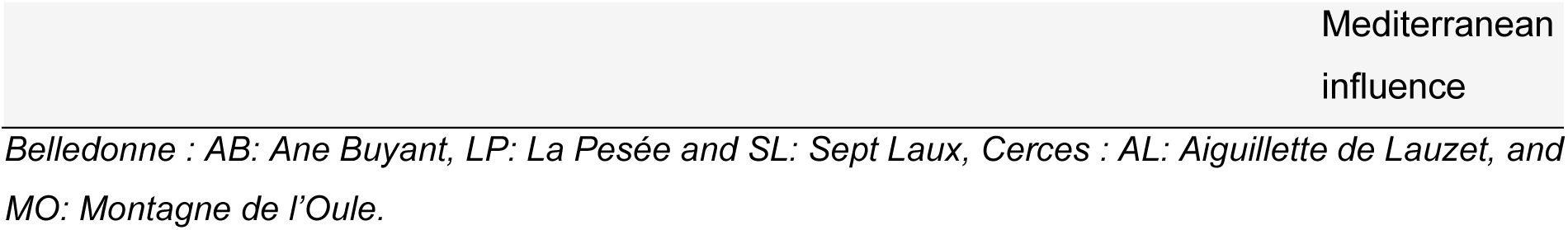
Descriptive data on the three mountain areas in the French Alps.

In the Champsaur mountain, the distribution of ibex range between 1320m and 3550m over 280km². Population size is estimated at 420 individuals. In Champsaur, the sheep herd includes 4 breeding farms for a total of 1070 sheep and 5 goats (Table 1).

Each sheep herd belongs to one farmer while several farmers grouped their sheep herds in Cerces and Champsaur (Table 1).

### Sample collection

We collected sheep feces for 15 days after their arrival on pasture to ensure that we collected nematode species representative of sheep at the time of arrival on pastures and not those ingested secondarily on alpine pastures. Similarly, ibex feces were collected prior to the arrival sheep and until 15 days after arrival to ensure that the nematode community was not influenced by the arrival of domestic livestock. We collected fresh ibex feces mostly directly on the ground and, in Belledonne, also during captures as part of the long-term monitoring program conducted by the French Office for Biodiversity. Ibex feces were collected within each of the pastoral units where we collected sheep feces. Whenever possible, feces were collected immediately after observation of ibex to avoid collecting of feces from the same individual. Feces from all age groups were collected. Samples were stored in plastic bags, sealed after air removal, and analyzed within 48h upon receipt in the parasitology laboratory of the National Veterinary School of Lyon (ENVL, Marcy-l’Étoile, France) or up to a maximum of 15 days after field collection (mean: 2.5 days). In total, we sampled 167 fecal samples from ibex and 90 fecal samples from 6 sheep herds, distributed over 6 pastoral units, i.e., Aiguillette du Lauzet, Montagne de l’Oule, Champsaur, Ane Buyant, La Pesée and Sept Laux (Table 2, Figure S1). In Belledonne, 21 samples were collected in 2018 following the sampling strategy of 2019. We controlled that year of sampling did not result in a significant change in the nemabiome of ibex (Figure S6) and included those samples in the analyses.

**Table 2:**
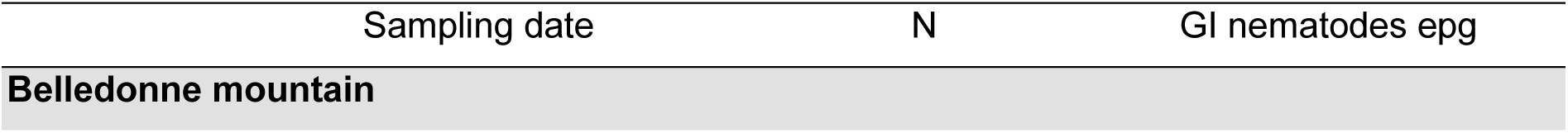
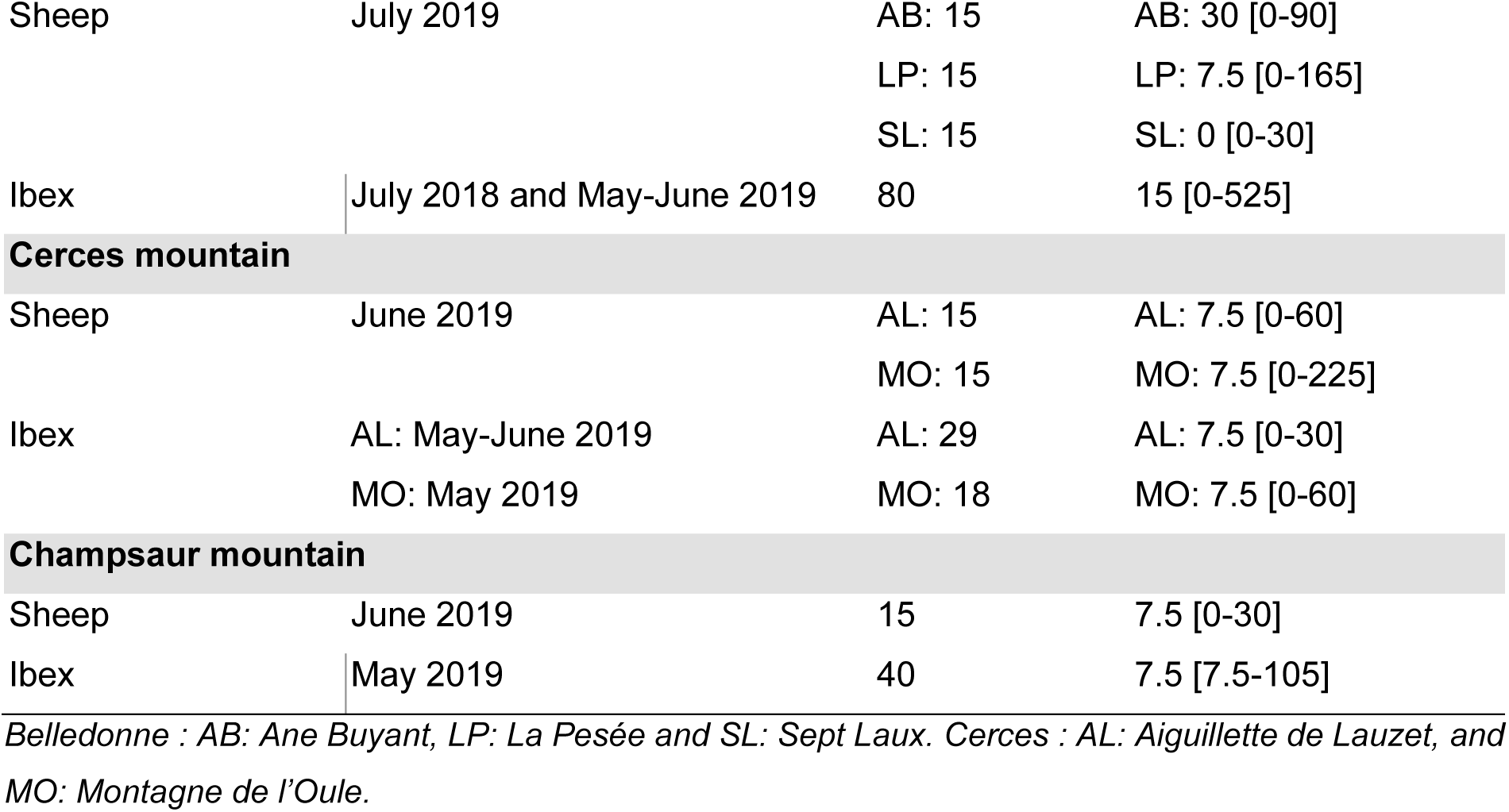
Sampling period and number of ibex and sheep feces collected in the three mountain areas of the French Alps. Coproscopic results of the gastro-intestinal (GI) nematodes (strongyles) in eggs per gram (epg) of feces are also reported (median[min-max]).

|  |  |  |  |
| --- | --- | --- | --- |
| Sheep | July 2019 | AB: 15<br>LP: 15<br>SL: 15 | AB: 30 [0-90]<br>LP: 7.5 [0-165]<br>SL: 0 [0-30] |
| Ibex | July 2018 and May-June 2019 | 80 | 15 [0-525] |
| **Cerces mountain** |  |  |  |
| Sheep | June 2019 | AL: 15<br>MO: 15 | AL: 7.5 [0-60]<br>MO: 7.5 [0-225] |
| Ibex | AL: May-June 2019<br>MO: May 2019 | AL: 29<br>MO: 18 | AL: 7.5 [0-30]<br>MO: 7.5 [0-60] |
| **Champsaur mountain** |  |  |  |
| Sheep | June 2019 | 15 | 7.5 [0-30] |
| Ibex | May 2019 | 40 | 7.5 [7.5-105] |

*Belledonne : AB: Ane Buyant, LP: La Pesée and SL: Sept Laux. Cerces : AL: Aiguillette de Lauzet, and MO: Montagne de l'Oule.*

### Parasitological analyses

The number of gastro-intestinal nematode eggs per gram of feces (epg) was counted following a modified McMaster protocol (Raynaud et al., 1970) with a solution of zinc sulphate (ZnSO_4_, density = 1.36, 1/15 dilution). The eggs were counted on a McMaster slide with two chambers (theoretical sensitivity of 15 epg). We also checked for the presence of low abundant parasite propagules with a “control slide”. We prepared this control slide with a 14mL tube filled with the remaining solution until a meniscus is formed. We covered the tube with a coverslip and centrifuged the tube with a basket centrifuge (5 min at 1200 rpm) to help propagules to rise and stick on the coverslip. After centrifugation, the coverslip was transferred onto a microscope slide for microscopical observation. We attributed the value of 7.5 epg for parasites with no egg observed on the McMaster, but at least one egg observed on the control slide (for a similar procedure see Beaumelle et al, 2021).

In order for strongyles to reach the L3 stage, coprocultures of feces were done at 24 ± 1 °C during 12-15 days with regular mixing and moistening. We then collected the L3 in tape water with a Baermann apparatus. We extracted gastrointestinal nematode DNA from samples for which there were at least 20 L3 and we limited the extraction to ∼200 L3. DNA was extracted using an extraction kit (Qiagen DNeasy^®^ PowerSoil) following the manufacturer’s instruction with an elution volume of 50 μl of water. We extracted the DNA of 30 randomly chosen samples twice as internal extraction controls (Taberlet et al., 2018). We quantified the DNA concentration for all samples using a Qubit 2.0 fluorometer (Life Technologies) and homogenized DNA samples to a DNA concentration of 1ng/μl (DNA samples were not diluted if the DNA concentration was <1 ng/µl).

### High throughput sequencing analyses

To determine the nemabiome of sheep and ibex, we used a modified version of the protocol developed by Avramenko et al (2015). The ITS2 region of the nuclear rDNA was amplified using the primer pair NC1 (Forward -5’-ACGTCTGGTTCAGGGTTGTT-3’) and NC2 (Reverse-5’-TTAGTTTCTTTTCCTCCGCT-3’) with the following PCR conditions: 10µl of Applied Biosystems™ Master Mix AmpliTaq Gold™ 360, 5.84µl of molecular biology grade water, 0.16µl of Bovine Serum Albumin, 2μL of 5 μM mixed F and R primers form, 2 μL of DNA lysate. The PCR was performed under the following conditions: 10 min initial denaturation at 95°C; followed by 35 cycles of denaturation (30 s at 95°C), annealing (30 s at 54°C), and extension (1 min at 72°C); a final extension at 72°C for 7 min.

To detect the mutations responsible for the resistance of gastrointestinal nematodes to benzimidazole, we used a modified protocol of Avramenko et al., (2019). Using the same mix as previously described, we amplified the β-tubulin isotype 1 fragment comprising the codons at position 167, 198 and 200 with two pairs of primers in two independent PCRs. The PCR was performed under the following conditions: 10 min initial denaturation at 95°C; 40 cycles of denaturation (30 s at 95°C), annealing (30 s at 65°C), and extension (30 s at 72°C); a final extension at 72°C for 7 min. We targeted *Teladorsagia circumcincta* and *Trichostrongylus* spp. (Forward:5’-CGCATTCWCTTGGAGGAGG-3’ and Reverse: 5’-GTGAGYTTCAAWGTGCGGAAG-3’) and *Haemonchus contortus* (Forward:5’-CGCATTCYTTGGGAGGAGG-3’ and Reverse: 5’-GTGAGTTTYAAGGTGCGGAAG-3’) with the primers described by Avramenko et al (2019). All forward and reverse primers were tagged at 5’ in order that each sample had a unique combination of tagged primers.

In all PCRs, we added positive PCR controls (i.e., *Haemonchus contortus* and *Teladorsagia circumcincta* DNA extracts), negative PCR controls (distilled H_2_O) and negative DNA extraction controls. All samples (including controls) were tagged with unique barcode identifiers to allow pooling into a single amplicon library (Taberlet et al., 2018), and all samples were independently amplified 4 times to ensure sequencing reliability. Amplifications were carried out in 96-well plates, totaling 209 ibex and sheep samples, 17 PCR positive controls, 13 PCR negative controls, 7 extraction negative controls, 30 DNA extraction controls, as well as 12 empty wells in each plate to quantify tag jumping during PCR and sequencing steps (Figure S2; De Barba et al., 2014; Taberlet et al., 2018).

All PCR products of the ITS2 and the two β-tubulin isotype 1 sets were purified using QIAquick® Spin Columns (QIAquick® PCR Purification KitQIAGEN) and quantified using a Qubit 2.0 fluorometer (Life Technologies). Next, we pooled the 3 purified DNA pools (ITS2, two β-tubulin isotype 1) based on their initial concentration and in proportion according to the following ratio: ITS2 50%, β-tubulin isotype 1 25% for each. According to preliminary tests, we expected to achieve a sequencing depth of 20 000 reads per ITS2 DNA sample and 5 000 reads per β-tubulin isotype 1 DNA sample. Sequencing was performed with pair-end sequencing technology on the Illumina platform (2*250 bp Miseq) at Fasteris, Geneva, Switzerland.

### Sequence analysis and taxon assignation

The sequence reads were first analyzed with the OBITOOLS package (Boyer et al., 2016). Forward and reverse reads were assembled with the *alignpairedend* function, and we kept only sequences with a good score of alignment (rnorm>0.8). Sequences were attributed to their samples with the *ngsfilter* function with default parameters. Subsequently, assigned sequences were analyzed with the dada2 package (Callahan et al., 2016) following the pipeline available in www.nemabiome.ca. The *dada2* pipeline returns Amplicon Sequence Variants (ASV) which are sequence variants differing by as little as one nucleotide (Callahan et al., 2017). Following Beaumelle et al. (2021), gastrointestinal nematodes were identified with four different methods of assignation-databases: BLASTn (Altschul et al., 1990) based on (1) the NCBI database (Accessed: November 2022), and (2) AssignTaxonomy (Callahan et al., 2016; Wang et al., 2007) and (3) IDTaxa (Murali et al., 2018) based on the nematode ITS2 rDNA database 1.1.0 (Workentine et al., 2020). To identify the species associated with the β-tubulin sequences, we used IDTaxa against the nematode β-tubulin isotype 1 DNA reference sequences supplied in Avramenko et al., (2019). We chose to attribute a confidence level to taxonomic identifications at the species level: high or moderate confidence if three or two methods of assignation, respectively, were congruent. We also adjusted the sequence filtering based on an adapted procedure of Calderón-Sanou et al. (2020). We kept only ASVs present in at least 2 replicates of the same sample and removed ASVs that were not assigned to the genus level for the ITS2 and to the species level for β-tubulin isotype 1. We removed potential contaminants (reagent contaminants and cross-contaminations) following the procedure detailed in Calderón-Sanou et al., (2020). For each sample, we summed the reads of the two replicates with the highest similarity (i.e., 1 -Bray-Curtis dissimilarity). If this similarity was lower than the mean similarity among all replicates, sample was discarded. Then, we verified that the 30 DNA extraction replicates had similar nemabiomes and kept one sample replicate out of two. At the end, we removed samples if they had <1000 reads of ITS2 and <500 reads of β-tubulin isotype 1 (Figure S3).

### Identification of non-synonymous mutations in codons 167, 198 and 200

For each nematode species, all β-tubulin isotype 1 ASVs were aligned to one of the β-tubulin isotype 1 consensus sequences of the reference database (Avramenko et al., 2019) using the *AlignSeqs* function of the DECIPHER package (Wright, 2016). We examined each β-tubulin isotype 1 ASV at codons 167,198 and 200 to determine if these codons were associated with non-synonymous mutations. However, we did not consider other polymorphic sites of the exon.

### Statistical analyses on measures of nemabiomes

We measured differences of nemabiomes between the two host species (sheep and ibex) across the 4 study sites (Aiguillette de Lauzet, Montagne de l’Oule, Champsaur and Belledonne). In the Cerces Massif, we considered independently the two sectors of the Aiguillette de Lauzet and the Montagne de l’Oule that correspond to distinct and distant pastures where sheep herds never met and where ibex feces were independently collected. We considered two measures of diversity using the abundance of ASVs, i.e., the alpha diversity and the beta diversity. The abundance of each ASV was computed as the relative frequency of reads for each ASV in each sample, i.e., read relative abundance (RRA). The alpha diversity was measured with the Shannon index that considers richness and evenness of communities, and the beta diversity was measured with the weighted UniFrac index estimated using the R phyloseq package (McMurdie and Holmes, 2013). The weighted UniFrac distance is a phylogenetic distance between the set of ASVs from each nemabiome weighted by the transformed abundance of each ASV (Lozupone and Knight, 2005). The phylogenetic distances were computed from a phylogenetic tree which was constructed using maximum likelihood with the GTR+G+I model according to the *ModelTest* function (Posada and Crandall, 1998; Schliep, 2011). The exact counts of ITS2 reads were transformed with the Hellinger transformation (e.g., square root of relative frequencies) to account for the high number of zeros in the community tables and to decrease the influence of rare ASVs in statistical analyses (Legendre and Legendre, 2012).

We tested the effects of host species and site, including their interaction, on alpha diversity with linear models, and on beta diversity using perMANOVA (*adonis2*, vegan R package (Oksanen et al., 2020)). All possible models including the null model were computed. For perMANOVA models, we used a custom function to compute Akaike’s information criterion corrected for small sample size (AICc) based on residual sums of squares (Dyson, 2018). In a model selection approach, for both alpha and beta diversity, all possible models were ranked using the AICc and we selected the model with the lowest AICc value. Models with ΔAICc ≤ 2 were considered equivalent (Burnham and Anderson, 2002), and in this case, we considered the most parsimonious one, i.e., the model with the lowest degrees of freedom.

As sex and age were determined for some ibex, we also tested the effect of ibex classes (1: adult males, 2: females or kids/yearlings) on alpha and beta diversity following the same model selection approach. This included ibex classes and sites as explanatory variables.

All analyses were carried out using R 3.6 (R Core team, 2020. https://www.R-project.org/).

### Statistical analyses on measures of resistant nematode strains

To compare the frequency of benzimidazole resistance in gastrointestinal nematodes between ibex and sheep, and among the 4 study sites, we tested whether host species, site and nematode species influenced the relative abundance of ASVs with a resistant allele. For this purpose, we used a generalized linear model with a binomial family and a model selection approach as described above.

We used *AlignSeqs* (Wright, 2016) to generate multi-sequence aligned β-tubulin isotype 1 haplotype data. For each gastrointestinal species, we removed short ASVs, e.g., ASVs with a sequence length <10% compared to the median ASV length. PopART v1.7 (Leigh and Bryant, 2015) was used to draw median joining networks based on the haplotype data of each gastrointestinal nematode species.

## Results

### Parasite material

The median number of eggs of strongyles per gram of feces were lower in sheep (7.5[0,148]_95%IQR_; n = 90) than in ibex (15[0,163]_95%IQR_; n = 167) feces (Mann–Whitney U test; *W* = 9043.5, *P* = 0.006). Due to the level of infestation in some samples, the number of L3 hatched from eggs was not sufficient (n<20) for 48 samples. These samples were not used for subsequent genetic investigations. Specifically, all samples from the Sept Laux sheep herd (n=15, Belledonne) were discarded. Therefore, the nemabiome was determined based on the ITS2 for 196 (n= 55 sheep and n=141 ibex) out of 209 samples for which DNA was extracted (Figure S1).

### Diversity of gastrointestinal nematodes in sheep and ibex

In total, we detected 408 ASVs corresponding to 13 gastrointestinal nematode species (Table 3, Figure S4). Eight ASVs were assigned to the genus level (i.e., *Marshallagia* spp., *Nematodirus* spp. and *Trichostrongylus* spp.) due to non-identical assignation among taxonomic methods. An ASV corresponding to the lungworm *Cystocaulus ocreatus* was discarded for the statistical analyses because we only focus on gastrointestinal nematodes.

**Table 3:** Number of ASVs, reads and samples for each nematode taxon, including the results from the ITS2 (nemabiome) and the β-tubulin isotype 1 (resistance to benzimidazole). N= number (and percentage) of samples in which a taxon was detected. Host species and study sites are mixed here.

| | ITS2 | | | β-tubulin isotype 1 | | |
| --- | --- | --- | --- | --- | --- | --- |
|  | ASVs | Reads | N (%) | ASVs | Reads | N (%) |
| *Bunostomum trigonocephalum* | 8 | 61 631 | 12 (6%) | - | - | - |
| *Chabertia ovina* | 19 | 95 585 | 62 (32%) | - | - | - |
| *Cooperia curticei* | 4 | 803 | 6 (3%) | - | - | - |
| *Cooperia fuelleborni* | 2 | 358 | 5 (3%) | - | - | - |
| *Haemonchus contortus* | 47 | 1 937 587 | 110 (56%) | 38 | 392 110 | 96 (62%) |
| *Marshallagia marshalli* | 23 | 338 064 | 57 (29%) | - | - | - |
| *Marshallagia* spp. | 2 | 404 | 2 (1%) | - | - | - |
| *Oesophagostomum venulosum* | 13 | 311 530 | 82 (42%) | - | - | - |
| *Ostertagia leptospicularis* | 1 | 5 | 1 (1%) | - | - | - |
| *Ostertagia ostertagi* | 2 | 269 | 2 (1%) | - | - | - |
| *Teladorsagia circumcincta* | 235 | 3 311 094 | 166 (85%) | 310 | 343 551 | 145 (94%) |
| *Trichostrongylus axei* | 26 | 462 936 | 103 (53%) | 33 | 68 414 | 104 (68%) |
| *Trichostrongylus colubriformis* | 12 | 411 776 | 102 (52%) | 8 | 144 643 | 112 (73%) |
| *Trichostrongylus vitrinus* | 9 | 521 817 | 123 (63%) | 44 | 92 834 | 107 (69%) |
| *Trichostrongylus* spp. | 5 | 71 124 | 70 (36%) | - | - | - |

*Teladorsagia circumcincta* was the most prevalent nematode species and was detected in 85% of samples (90%, n=127/141 ibex and 71%, n=39/55 sheep), followed by *Trichostrongylus vitrinus,* 63% (73%, n=103/141 ibex and 36%, n=20/55 sheep) and *Haemonchus contortus,* 56% (70%, n=98/141 ibex and 22%, n=12/55 sheep) (Table 3, Figure S4). *Nematodirus* spp. and *Ostertagia leptospicularis* were the rarest species and were detected in only 2 and 1 sample, respectively and with a very low relative frequency (<0.1%).

The parasites of the genus *Nematodirus* were not considered for the following results as our coproculture protocol is not the most appropriate for these parasites (i.e., some species need more than 2 weeks to reach the L3 stage and need to be exposed to cold temperatures before coproculture, which can be deleterious for other strongyle species such as *Haemonchus contortus*, van Wyk and Mayhew, 2013).

According to the model selection approach, host species was the only factor that explained the alpha diversity (Table S1). The model indicated that sheep had a lower alpha diversity compared to ibex (β = −0.42 ± 0.07, *P* < 0.001, R² of the model=0.15) (Figure 2b). Site, and its interaction with host species were not retained in the selected model (Table S1).

**Figure 2:**
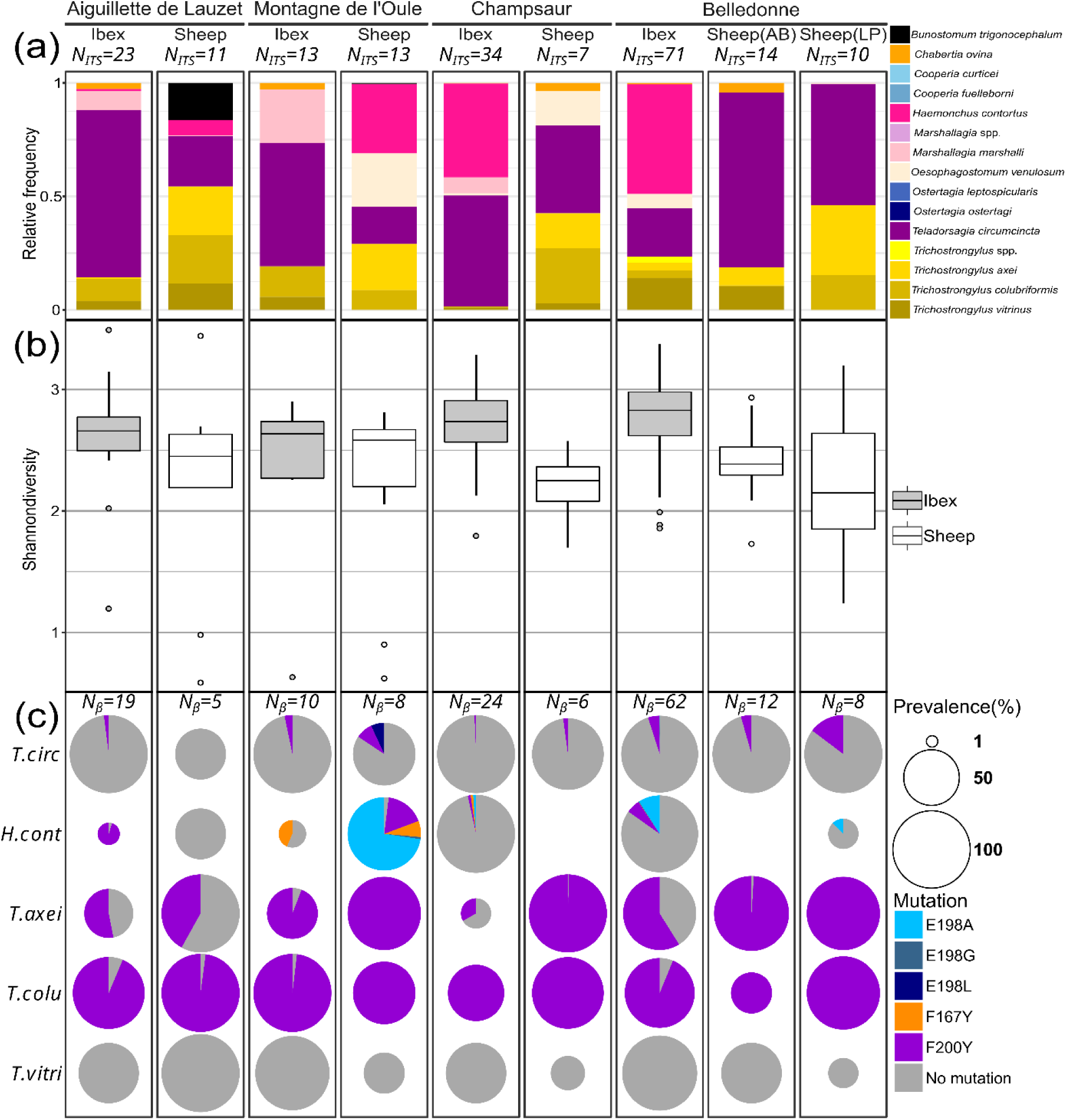
(a) Mean relative frequencies of gastrointestinal nematode species, (b) Shannon diversity of ITS2 ASVs and (c) prevalence and mean relative frequencies of β-tubulin isotype 1. Results and sample size (N_ITS_ and N_β_) are presented for each host species (sheep or ibex) in each study site (Cerces: Montagne de l’Oule and Aiguillette de Lauzet; Champsaur and Belledonne: Ane Buyant (AB) and La Pesée (LP)). On panel (c), the size of the pie chart corresponds to the prevalence of the corresponding gastrointestinal nematode species in the population, and the size of each slice to the mean proportion of each allele.

Beta diversity was best explained by both factors, host species (F_1,188_ = 27.69, *P* = 0.001) and site (F_3,188_ = 16.39, *P* = 0.001), and their interaction (F_3,188_ = 25.61, *P* = 0.001) according to model selection results (Table 4, Table S2). Some gastrointestinal nematodes were mostly found in sheep feces, or only in ibex feces (Figure 2a). *Trichostrongylus colubriformis* was more frequent in the Ecrins National Park, i.e., in the Cerces and Champsaur massifs combined (mean RRA: 10% [6%; 13%]_95CI_) than in Belledonne (mean RRA: 4% [2%; 6%]_95CI_). Likewise, *Marshallagia* spp. was more frequent in ibex feces in Cerces and Champsaur mountains (mean RRA: 11% [6%;15%]_95CI_) than in ibex feces in Belledonne (0.4% [-0.2%; 1%]_95CI_). The distribution of *Haemonchus contortus* in host species and sites had a particular pattern. This parasite was more frequent in ibex feces compared to sheep feces in Belledonne (mean RRA: 0.004% in sheep; 48% in ibex) and Champsaur (mean RRA: 0% in sheep; 41% in ibex), while the opposite was observed in the Cerces (Aiguillette du Lauzet: mean RRA of 7% in sheep and 0.9% in ibex; Montagne de l’Oule: mean RRA of 30% in sheep and 0% in ibex).

**Table 4:** Parameters estimated from the best PerMANOVA model explaining the beta diversity in ibex and sheep. The effect of host species (ibex or sheep) and site (Aiguillette de Lauzet, Montagne de l’Oule, Champsaur or Belledonne) and the interaction between the two factors are reported. Partial R² are reported with the corresponding *F-*value and *p value* (P).

| Diversity index | Best model selected | Variables | partial $R^2$ | F-value | P |
| --- | --- | --- | --- | --- | --- |
| Weighted UniFrac | $\beta \sim \text{Site} \times \text{Host species}$ | Residuals | 0.55 | - | - |
|  |  | Site | 0.14 | 16.39 | 0.001 |
|  |  | Host species | 0.08 | 27.69 | 0.001 |
|  |  | Site: Host species | 0.22 | 25.61 | 0.001 |

Concerning ibex only, the model selection approach retained the effects of site and class of individuals and the interaction between the two independent factors in the best model explaining alpha diversity (Table S3). We found significant differences among sites: Shannon index of alpha diversity was higher in Belledonne (β = 0.62 ± 0.16, P < 0.001, R² of the model=0.27) and Champsaur (β = 0.55 ± 0.16, P < 0.001) compared to the Aiguillette du Lauzet. The diversity of nematodes was higher also in males compared to females/yearlings (β = 0.60 ± 0.15, P < 0.001), except in Champsaur where males presented a lower alpha diversity than females with kids (β = −0.52 ± 0.20, P < 0.001). For the beta diversity, the best model included only the effect of site on the difference of nemabiomes among ibex (F_2,62_ = 15.93, P = 0.001, Table S4).

### Anthelmintic resistance

We found 433 different β-tubulin isotype 1 ASVs in 154 (n= 39 sheep and n=115 ibex) out of 209 samples for which DNA was extracted. Among the 5 gastrointestinal nematode species targeted by specific primers, we detected *Haemonchus contortus* in 96 samples, *Teladorsagia circumcincta* in 145 samples, *Trichostrongylus axei* in 104 samples, *Trichostrongylus vitrinus* in 107 samples and *Trichostrongylus colubriformis* in 112 samples (Table 3, Figure S5). No resistance mutation was detected for *Trichostrongylus vitrinus.* Therefore, *Trichostrongylus vitrinus* was not included in the model explaining the relative abundance of resistant reads.

Resistance mutations were highly frequent (93.5%; n=144/154) with only 10 ibex feces (3 from Belledonne and 7 from Champsaur) in which no resistant mutation was detected. Based on the best model for resistant RRA, the frequency of resistant nematodes depended on gastrointestinal nematode species and the interaction between host species and the study site (Table 5, Table S5). *Teladorsagia circumcincta* was the species with the lowest resistant RRA and *Trichostrongylus colubriformis* had the highest resistant RRA (Table 5). The mean observed RRA of resistant nematodes differed between gastrointestinal nematode species (*Haemonchus contortus:* 19% [13;25]_95CI_*; Teladorsagia circumcincta:* 4% [3;6]_95CI_; *Trichostrongylus axei:* 70% [63;78]_95CI_; *Trichostrongylus colubriformis:* 96% [93;99]_95CI_). Resistant RRA were generally lower in ibex compared to sheep (β = −2.21 ± 0.66, *P* < 0.001). Resistant RRA were the lowest in the Aiguillette de Lauzet but this was also the only site where ibex had a significantly higher resistant RRA compared to sheep (Table 5). We found no significative effect of ibex classes (males or females and kids/yearlings) on benzimidazole resistance frequencies (*F*_1,49_=0.03, P = 0.863, ANOVA test).

**Table 5:** Parameter estimates for the best generalized linear model explaining the resistant reads relative abundance (RRA) in ibex and sheep. The effect of host species (sheep as reference), study sites (Belledonne as reference), and their interaction, in addition to the nematode species (*Teladorsagia circumcincta* as reference) are reported. Parameter estimates with standard error (SE) are reported with the corresponding *z-value (z*-val) and *p value* (*P*). AL: Aiguillette de Lauzet; MO: Montagne de l’Oule; Ch: Champsaur; Hc : *Haemonchus contortus*; Ta: *Trichostrongylus axei*; Tcol: *Trichostrongylus colubriformis*.

| Best model selected | Variables | Parameter estimate ± SE | z-val | P |
| --- | --- | --- | --- | --- |
| Resistant RRA ~ host species x study sites + nematode species | Intercept | -2.02 ± 0.59 | -3.45 | **5e-04** |
| | Species | -2.21 ± 0.66 | -3.35 | **8e-04** |
| | Mountain(AL) | -2.89 ± 0.99 | -2.93 | **0.003** |
| | Mountain(MO) | -1.37 ± 0.89 | 1.54 | 0.123 |
| | Mountain(Ch) | -0.10 ± 1.18 | -0.08 | 0.935 |
| | Nematode(Hc) | 2.52 ± 0.62 | 4.05 | **5e-05** |
| | Nematode(Ta) | 4.68 ± 0.59 | 7.95 | **1e-15** |
| | Nematode(Tcol) | 7.38 ± 0.75 | 9.79 | **<2e-16** |
| | Species:Site(AL) | 2.90 ± 1.19 | 2.44 | **0.015** |
| | Species:Site(MO) | 0.04 ± 1.28 | 0.03 | 0.973 |
| | Species:Site(Ch) | -0.75 ± 1.32 | -0.56 | 0.574 |

The most frequent resistance mutation was the F200Y, present in the 4 gastrointestinal nematode species and 141 fecal samples, followed by the E198A (46 samples, 2 nematode species: *Haemonchus contortus* and *Teladorsagia circumcincta*), the F167Y (17 samples, 2 nematode species: *Haemonchus contortus* and *Teladorsagia circumcincta*), the E198L (7 samples, *Teladorsagia circumcincta*) and the E198G (4 samples, *Haemonchus contortus*) (Figure 2c).

Sheep and ibex shared 164 (38%) β-tubulin isotype 1 haplotypes and 238 (55%) β-tubulin isotype 1 haplotypes were only found in ibex samples (Figure 3). Most of the resistant haplotypes of *Teladorsagia circumcincta* and *Haemonchus contortus*, e.g., containing a non-synonymous mutation at the codon 167, 198 or 200, were genetic variants of a common sensitive haplotype shared by ibex and sheep (Figure 3). The resistant haplotypes of *Trichostrongylus axei* and *Trichostrongylus colubriformis* were more common than the sensitive haplotypes and the most similar sensitive haplotypes were found either in both sheep and ibex samples, or only in ibex samples. Both, *Trichostrongylus colubriformis* and *Trichostrongylus vitrinus* showed two distinct lineages, separated by ≥10 mutations. One of the lineages of *Trichostrongylus vitrinus* was only found in ibex from Belledonne and an ibex from Champsaur (Figure 3) while the ASVs of the second lineage were found both in ibex and sheep. One of the lineages of the *Teladorsagia circumcincta* was more diverse (Figure 3).

**Figure 3:**
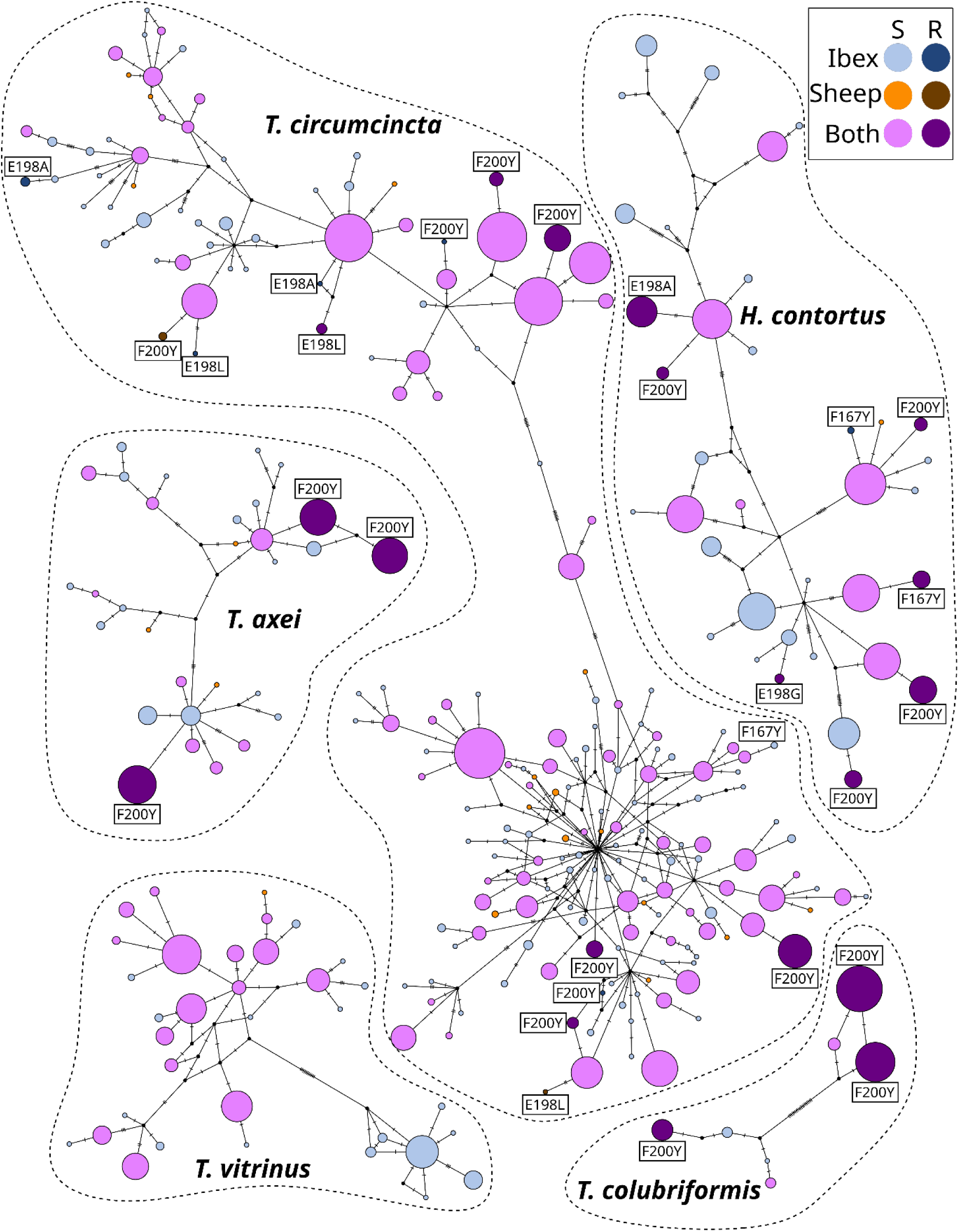
Median joining network of β-tubulin isotype 1 haplotypes. Each point represents a unique haplotype, and the colors correspond to the host species in which the haplotype was detected. The size of the point is proportional to the number of samples in which the haplotype was found. S: sensitive haplotype, R: resistant haplotype. The tag above the points indicates the name of the mutation, based on the codon position and the substitution of the amino acid.

## Discussion

Because resident Alpine ibex use pastures grazed by transhumant sheep during the summer, but not concurrently, we sought to assess the extent of nematode sharing between these two host species. Specifically, we investigated the presence of anthelmintic-resistant nematode strains in sheep and ibex to determine the role of transhumant sheep in contaminating alpine pastures, and whether ibex may contribute to the circulation and maintenance of anthelmintic resistant nematodes. We used a metabarcoding approach based on the sequencing of ITS2 and β-tubulin to demonstrate that both sheep and ibex were infected by the same gastrointestinal nematode species and shared anthelmintic-resistant strains, despite the absence of sheep on alpine pastures for much of the year and therefore a narrow temporal window for contamination.

In line with other studies investigating the gastrointestinal nematodes of sheep and ibex (Burgess et al., 2012; Gruner et al., 2006; Redman et al., 2019; Zaffaroni et al., 2000), the most prevalent and abundant species in both host species was *Teladorsagia circumcincta*. Next, *Trichostrongylus vitrinus* was moderately prevalent, but not abundant in sheep or ibex nemabiomes. These results are consistent with the year-round climatic conditions as these two nematode species, along with *Marshallagia* spp., are better adapted to cold temperatures than the other nematode species detected in this study (O’Connor et al., 2006; Zaffaroni et al., 2000). The studied sheep flocks originate from the French plains and/or the south of France and are driven into mountain areas in early summer. Consequently, their nemabiome at the time of sampling is representative of the gastrointestinal nematode communities present in sheep on the farm, prior to transhumance. In a similar context, Gruner et al., (2006) observed a high prevalence of *Teladorsagia circumcincta* in two of three transhumant sheep flocks at the beginning of the grazing season in the southern Alps. Furthermore, transhumant sheep flocks appear to ingest mainly *Teladorsagia circumcincta* when grazing in the mountains, as this parasite remains the dominant species identified in feces and tracer lambs during the summer (Gruner et al., 2006). Pastoral activity in mountainous areas of France could therefore favor nematode species more adapted to cool and wet environmental conditions, such as *Teladorsagia circumcincta* (O’Connor et al., 2006), compared with sheep grazing on the plains year round. To confirm this hypothesis, the nemabiome of transhumant sheep should be compared with the nemabiome of resident sheep that stay in farm all year around.

A high relative frequency (>30%) of *Haemonchus contortus* was detected in ibex in Belledonne and Champsaur. In contrast, almost no *Haemonchus contortus* was observed in sheep flocks driven to these mountains, raising the possibility that ibex may be contributing to the infection of sheep with *Haemonchus contortus*. To our knowledge, this is the first time that such a relative abundance of *Haemonchus contortus* is reported in Alpine ibex (see previous studies based on morphological identification: Carcereri et al., 2021; Marreros et al., 2012; Zaffaroni et al., 2000). In addition, it should be noted that Alpine ibex were sampled before potential contamination by domestic sheep could be detected, i.e., before the end of the pre-patent period (i.e., the time between infection and egg production) and at the end of spring – to early summer, which is the start of the epidemiological period for *Haemonchus contortus* infection in high-altitude mountain areas. We can therefore expect higher levels of contamination in late summer, when domestic sheep leave mountain pastures. Nonetheless, we cannot exclude the possibility that some laboratory issues might have reduced the apparent prevalence and abundance of *Haemonchus contortus*. For instance, some samples from sheep were kept in the fridge at 4°C during 2 to 3 days (including the sheep samples without *Haemonchus contortus* from Belledonne and Champsaur) which could have reduced the proportion of *Haemonchus contortus* eggs hatching (McKenna, 1998).

The detection of *Haemonchus contortus* raises conservation issues for Alpine ibex as this nematode species is known to be highly pathogenic in sheep (Taylor et al., 2015). Infection of a phylogenetically related species, the Pyrenean ibex (*Capra pyrenaica pyrenaica*), with a few thousand *Haemonchus contortus* resulted in severe clinical signs, including extremely low weight and hemorrhagic anemia (Lavín et al., 1997). Additionally, *Haemonchus contortus* may have contributed to the collapse of the Northern Chamois, *Rupicapra rupicapra*, in the province of Lecco, Italy from November 2000 to March 2001 (Citterio et al., 2006). As gastrointestinal nematodes can have an impact on the demographic dynamics of host populations (Acerini et al., 2022; Albery et al., 2021; Albon et al., 2002), they are suspected of being behind the low natality rates observed in the French Alpine ibex populations (Brambilla et al., 2020). While Alpine ibex appear to be fairly resilient to parasite infections (Marreros et al., 2012), further investigations should be carried out to assess the consequences of gastrointestinal nematode infections for ibex at both individual and population levels.

Several of the nematode species we found are common to other ungulate species present in the study areas (Mediterranean mouflon, *Ovis gmelini musimon × Ovis sp*.; Northen chamois; domestic goat, *Capra hircus*; red deer*, Cervus elaphus*; and roe deer, *Capreolus capreolus*) (Zaffaroni et al., 2000). In our study, only a few domestic goats were present in Belledonne (n = 11 individuals) and in Champsaur (n =5 individuals) and represented less than 0.01% of the domestic flock in these areas. As we collected feces directly on the ground, we cannot exclude that goat feces were collected instead of sheep feces. In our opinion, domestic goats should not have a significative influence on nemabiome of ibex in our study area considering the scarcity of the species. In future analyses, we should consider the different domestic and wild ungulate species living in the same study area. Especially because they have different space use, different nemabiomes and could provide key information to better understand the dynamic of nematode exchanges among domestic and wild ungulates. We found a higher diversity of nematodes in adult males compared to females and kids/yearlings. Sexual parasitism towards males is commonly observed in vertebrates and ungulates in particular (Klein, 2000; Martínez-Guijosa et al., 2015; Oliver-Guimerá et al., 2017, but see Beaumelle et al., 2021; Bourgoin et al., 2021). This parasitism towards males is generally explained by both hormonal and behavioral differences between the two sexes. Generally, males tend to allocate more energy to the development of traits influenced by testosterone, such as secondary sexual characteristics (e.g. the length of horns in ungulates) or courtship displays (e.g. male aggression for mating opportunities). It is important to note that while testosterone is necessary for the development of these secondary sexual characteristics in males, high levels of testosterone have also been linked to an altered immune system (Klein, 2004), leading to increased parasitism. In addition, ibex segregate by sex (Brambilla et al., 2022), providing less opportunities for intersexual transmission of parasites. Contrary to females and kids, males feed on patches grazed by domestic sheep before the grazing period (Margaillan, 2021), increasing the probability of infection of males by over-wintering nematodes deposited by livestock during the previous transhumance (O’Connor et al., 2006). However, we did not observe any difference between classes concerning benzimidazole resistance frequencies. Further information on the spatial distribution of both sexes is necessary to investigate the susceptibility of different sexes or age classes and its importance in the spread and exchange of parasites with domestic livestock (Bourgoin et al., 2021).

We detected anthelmintic resistant alleles in 4 out of the 5 nematode species tested, namely *Haemonchus contortus*, *Teladorsagia circumcincta*, *Trichostrongylus axei*, *Trichostrongylus colubriformis*, but not in *Trichostrongylus vitrinus*. Both sheep and ibex hosted resistant strains of the 4 nematode species and only 10 out of 116 ibex carried only susceptible strains. Benzimidazole resistance was therefore very common in both host groups, consistent with the situation of sheep farms in Europe (Rose et al., 2015; Rose Vineer et al., 2020). The presence of anthelmintic resistant nematodes in ibex is most likely explained by the indirect transmission of resistant nematodes from sheep to ibex through the deposition of infected feces by sheep on pastures. The large number of shared β-tubulin ASVs between sheep and ibex and the high overlap between their nemabiomes tend to confirm this hypothesis (Figure 2c, Figure 3). This is also in accordance with other studies investigating nematode parasite exchange at the interface between wild and domestic ungulates (Beaumelle et al., 2022; Cerutti et al., 2010; Laca Megyesi et al., 2019). Another pathway for the contamination of ibex by resistant nematodes could be the sublethal exposure of the free-living stages of nematodes to anthelmintic residues excreted in the environment by treated sheep, which can select for anthelmintic resistance in situ (Dimunová et al., 2022). Whereas sheep are generally treated just before their ascent to the mountain pastures, excretion of anthelmintic drugs via sheep feces can occur during several days after the administration and molecule degradation can last days, or even months (Kolar et al., 2006). Unfortunately, the level of drugs in the environment, their persistence, and their spread in grazed mountainous area are entirely unknown. Environmental circulation of anthelmintic residues should be investigated in further studies to understand their impact on the presence of resistant nematodes in wildlife.

It is worth noting that feces of ibex were sampled before the arrival of sheep on pastures. This demonstrates that anthelmintic resistant nematodes can be maintained in mountainous areas from year to year in wild populations of ibex, despite harsh winter environmental conditions, and in the absence of the main source of parasites during most of the year, i.e., domestic sheep. The shedding of eggs from resistant nematodes by ibex prior to the arrival of domestic sheep suggests the potential role of ibex as a reservoir of anthelmintic resistant nematodes for other susceptible domestic and wild ungulates. Once resistant strains have been selected, the absence of selection pressure (i.e. absence of the use of anthelmintics) does not guarantee the reversion of resistance (Hamilton et al., 2022; Leathwick et al., 2015). Consequently, ibex may maintain benzimidazole-resistant strains for several years even in the absence of selection pressure. In addition, the position of resistant mutant strains detected at the periphery of haplotype networks (Figure 3) supports the lack of benzimidazole resistant reversions and relatively recent selection of benzimidazole resistance. The recent selection of resistance could result from the repeated use of benzimidazole and possibly from the presence of anthelmintic drug residues in the environment which maintain a selection pressure for gastro-intestinal nematodes (Dimunová et al., 2022). The 5 nematode species studied seemed to have different selection dynamics which may reflect their life history traits (Redman et al., 2015). In fact, we detected no resistant allele in *Trichostrongylus vitrinus* and conversely, the proportion of benzimidazole resistant strains of *Trichostrongylus axei* and *Trichostrongylus colubriformis* were high in sheep, i.e., between 40% and 100% and slightly lower in ibex, i.e., 33-100% (Figure 2c). These results are consistent with observations in the literature. Strains of *Trichostrongylus vitrinus* resistant to benzimidazoles were also rare in other studies of sheep farms, e.g., in the UK (Avramenko et al., 2019), and in Canada (Queiroz et al., 2020). Conversely, higher frequencies of benzimidazole resistance in *Trichostrongylus axei* and *Trichostrongylus colubriformis* were already reported in sheep, e.g., in UK (*Trichostrongylus axei*: 26-27% and *Trichostrongylus colubriformis*: 53-62%; Avramenko et al., 2019), in Austria (*Trichostrongylus colubriformis:* 77%-100%; Hinney et al., 2020), or in France, (*Trichostrongylus axei*: 63%; Palcy et al., 2010). Furthermore, we found a low proportion of resistance in *Teladorsagia circumcincta* both in sheep and ibex (mean ± standard error of the mean: 6.6 ± 3.5%), in comparison, for example, with *Trichostrongylus axei* and *Trichostrongylus colubriformis*, (Figure 2c). This contrast with previous results from the literature, where a higher frequency of resistance alleles was found for *Teladorsagia circumcincta*: 32.4 ± 6.8% in Hinney et al. (2020) and ∼66% in Avramenko et al. (2019). Regarding *Haemonchus contortus*, we found high variability of resistance frequencies among sheep flocks (69.9 ± 14.4%). Our results are similar to the observations of Avramenko et al. (2019), but contrast with those of Hinney et al. (2020), who detected high resistance frequencies for *Haemonchus contortus* (91.9 ± 3.7%) in transhumant sheep flocks in the Austrian Alps. In addition, *Haemonchus contortus* was the parasite species with the highest diversity in terms of non-synonymous mutations. Besides the commonly found polymorphisms at codons 167 and 200 (Avramenko et al., 2019; Redman et al., 2015), we detected several mutations at codon 198, occurring in particular with high frequency on the sheep farm in Montagne de l’Oule. In contrast to our study, previous research found polymorphisms at codon 198, but they were rare (Francis and Šlapeta, 2023; Ramünke et al., 2016).

Several factors are suspected to contribute to interspecific differences in the selection of resistance strains among nematode species, including specific reproductive rates, seasonal dynamics, exposure to anthelmintics in the digestive tract, drug effective dose and the fitness cost of benzimidazole resistance, and climatic conditions in the location of sheep farms, anthelmintic strategies, e.g., treatment molecules, timing and rate of anthelmintic treatments and grazing management (Hodgkinson et al., 2019; Redman et al., 2015). However, the links between parasite traits and interspecific variation in resistance acquisition by gastrointestinal nematodes has not been tested yet (Morgan et al., 2019). Our results suggest some clues related to the ecology of the nematode species.

Firstly, nematode species have different abilities to practice hypobiosis, i.e., the ability to halt embryonic development under environmental constraints (Gibbs, 1986). *Haemonchus contortus* and *Teladorsagia circumcincta* are known to arrest development more frequently than *Trichostrongylus* spp. (Langrová et al., 2008), and hypobiotic larvae have been shown to be less sensitive to drugs (Sargison et al., 2007). Secondly, among the *Trichostrongylus* spp., *Trichostrongylus vitrinus* may have a higher proportion of overwintering larvae in pastures as this species is more resistant to cold temperature compared to *Trichostrongylus axei* and *Trichostrongylus colubriformis* (O’Connor et al., 2006). As parasites on pastures may be less subject to selection pressure by anthelmintics, they could be a source of susceptible strains.

As the proportion of resistant strains is generally lower in ibex compared to sheep, ibex may contribute to a dilution effect of resistant strains, i.e., by hosting susceptible nematodes. However, the role of ibex in the maintenance of a refugia needs to be investigated by considering the relative number of susceptible strains deposited by ibex on a pasture compared with sheep. Furthermore, it seems that the role of ibex in the maintenance of a refugia may vary according to nematode species. For example, ibex excrete a lower proportion of *Trichostrongylus axei* eggs than sheep (Figure 2a), but these eggs largely contain resistant strains (Figure 2c). In contrast, *Teladorsagia circumcincta* and *Haemonchus contortus* in ibex were more frequently susceptible and genetically diverse (higher number of ITS2 and β-tubulin ASVs) compared with *Trichostrongylus axei* and *Trichostrongylus colubriformis* in ibex (Table 3, Figure 3). As *Teladorsagia circumcincta* was dominant in ibex, a refuge of susceptible *Teladorsagia circumcincta* strains may be maintained within ibex and may contribute to limiting the spread of resistance in sheep farms. Nematodes can negatively or positively interact within the host gut, and interactions between species or between strains may have important implications for the selection of resistance. However, the magnitude of within-host interactions between nematode strains/species and its implication in the management of resistance remains to be determined (Hellard et al., 2015; Lello et al., 2004).

Differences in patterns among massifs were observed at the community and genetic level among sheep flocks and ibex populations. Indeed, it was expected that differences in nemabiome composition would be observed between the massifs and sheep flocks, considering that sheep and ibex from different areas never meet (R. Papet, C. Toïgo, E. Vannard, Pers. communication). Furthermore, the sheep flocks come from different locations and have been subjected to different anthelmintic strategies. For the ibex, differences in the original population of translocated animals (Gauthier and Villaret, 1990; Kessler et al., 2022) and potential founder effects – not all parasites present in the source population were present in the newly established population - may have had a long term impact on the composition of the nemabiome. For example, the distinct population of *Trichostrongylus vitrinus*, found mainly in ibex in Belledonne, may have been inherited from the founding ibex population. This highlights that various reintroductions of ibex in the study area can also influence the composition of the parasite community. This distinct population *of Trichostrongylus vitrinus* was absent in sheep grazing in Belledonne, which hosted other strains of *Trichostrongylus vitrinus* despite summers of co-occurrence of sheep and ibex. It is possible that this strain is adapted to ibex and incapable of developing in sheep or that this strain is highly sensitive to antiparasitic treatments used in sheep farms and is systematically eliminated when sheep are treated. However, it is possible that this observation may also be due to a sampling bias as the number of sheep sampled remains low.

In conclusion, transmission of gastrointestinal nematode species, including resistant nematode strains, occurs between sheep and ibex even though the contact between the two species is limited to the summer period. Our study specifically demonstrates that ibex can maintain and shed eggs of resistant gastrointestinal nematodes even in the absence of sheep on pastures for several months, suggesting a potential reservoir role for ibex. However, the extent to which each host species can influence the nematode community of the other during the transhumant period remains to be determined. A temporal sampling before, during and after the different host species share the same pasture - should be considered for future studies. Additionally, analysis of parasite population structure using appropriate genetic markers such as microsatellites or SNPs could quantify gene flow between ibex and sheep nematode populations (Cerutti et al., 2010). Next, intervention studies could be employed to infer the role of ibex in maintaining nematode populations shared between the two host species (Viana et al., 2014). Experimental infections of captive ibex or monitoring free-ranging ibex populations after access to alpine pastures has been restricted to livestock should help us to refine the ability of ibex to maintain nematodes from domestic ungulates, including resistant nematodes. Finally, epidemiological models could be useful tools to better understand the dynamics of resistant parasites at the livestock-wildlife interface (Brown et al., 2022; Dickinson et al., 2024). The lower proportion of resistance alleles in ibex compared to sheep underlines the possibility that ibex could contribute to the maintenance and circulation of susceptible strains in sheep. As with roe deer (Beaumelle et al., 2022), domestic sheep contribute to the modification of the nemabiome of ibex. This raises concerns about ibex conservation, and the consequences of strongyle infection in ibex should be investigated. Indeed, ibex are characterized by low genetic diversity due to the severe demographic decline of this species followed by multiple re-introductions (Grossen et al., 2018). The high genetic structure of immunity-related loci among ibex populations (Kessler et al., 2022) raises additional concerns, as both neutral and adaptive genetic diversity are known to have an influence on parasite resistance and tolerance in ungulates (Portanier et al., 2019).

## Appendices

Supplementary data to this article can be found online in MendeleyData (DOI: 10.17632/cm97cg87d6.1).

## Acknowledgements

The authors warmly thank all the professionals from the Office Français de la Biodiversité and all the trainees for data collection, J. S. Gilleard and E. Redman from the University of Calgary, C. Lionnet from the Laboratoire d’Ecologie Alpine (LECA) and other members of the lab to help us developing the deep sequencing analyses for nematodes. The research benefited from the support of AnaBM (USMB) and AEEM (UGA) laboratory facilities. We warmly thank Karen McCoy as well as two anonymous reviewers for their insightful comments on the manuscript.

## Data, scripts, and supplementary information availability

The bioinformatic pipeline, the ASV analysed during the current study and the R script of statistic analyses are available in MendeleyData (DOI: 10.17632/cm97cg87d6.1).

