## Supplementary figures and tables for "Cross-transmission of resistant gastrointestinal nematodes between wildlife and transhumant sheep"

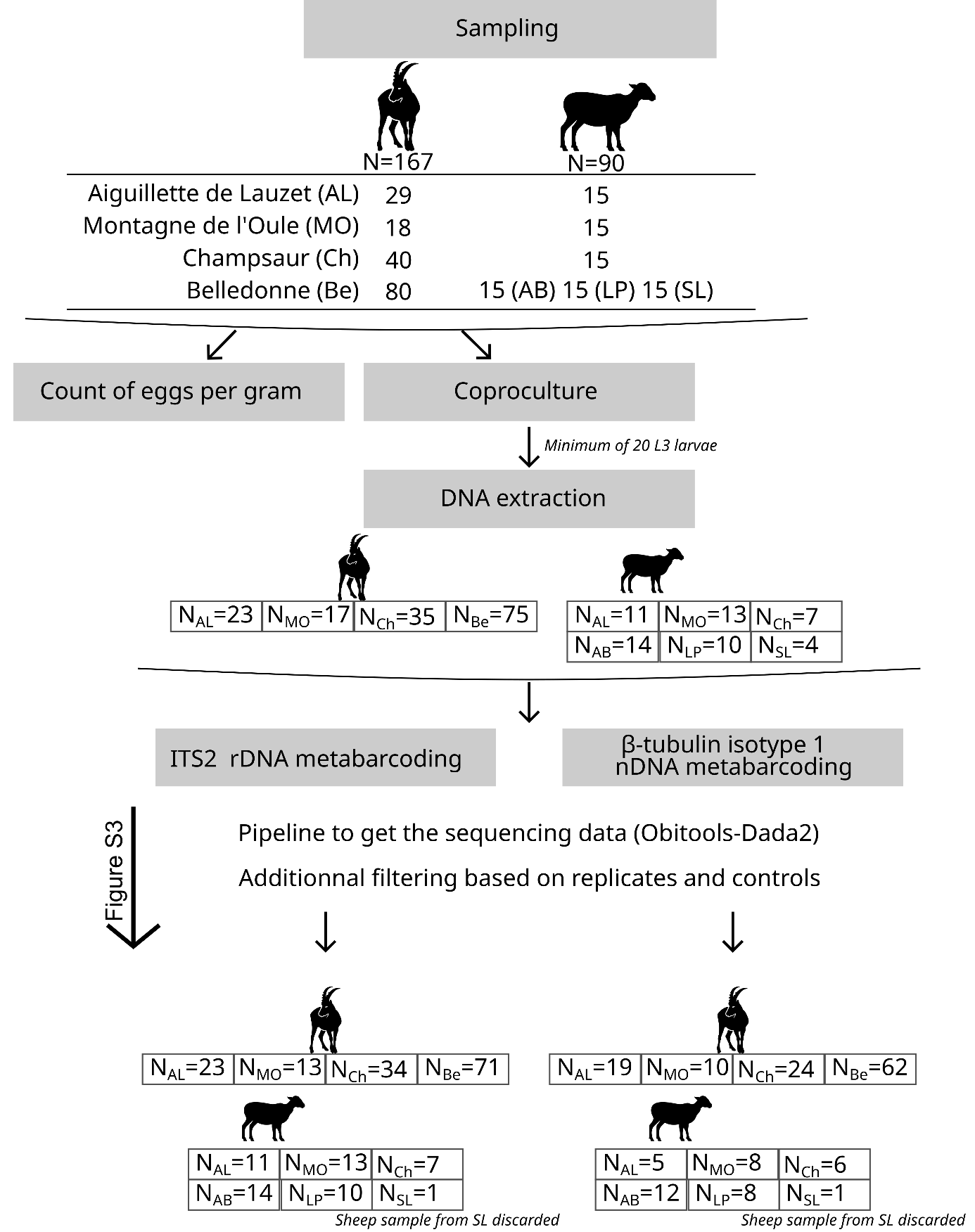

Figure S1. Workflow of the data analyses from the sampling to the results. AB: Ane Buyant, LP : La Pesée and SL : Sept Laux.

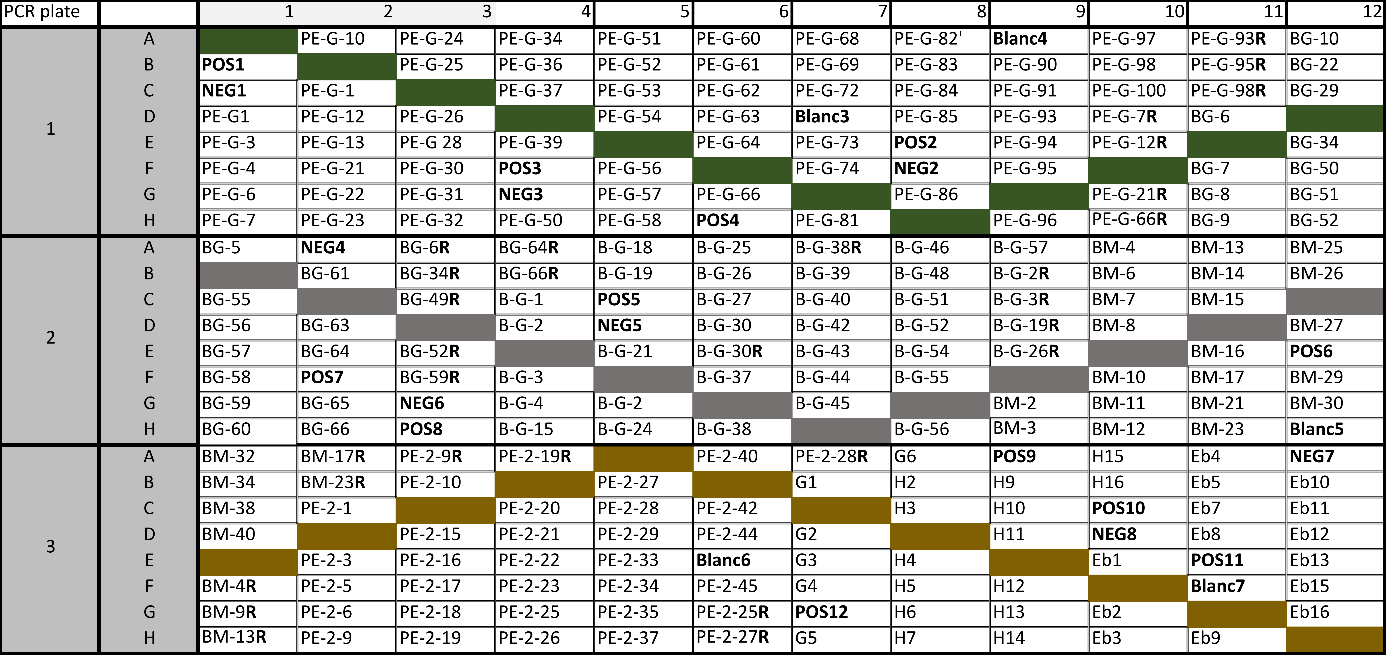

Figure S2. Position of samples in the PCR plate. POS: positive controls, NEG: negative controls, Blanc: negative controls of DNA extraction, R: DNA extraction replicates.

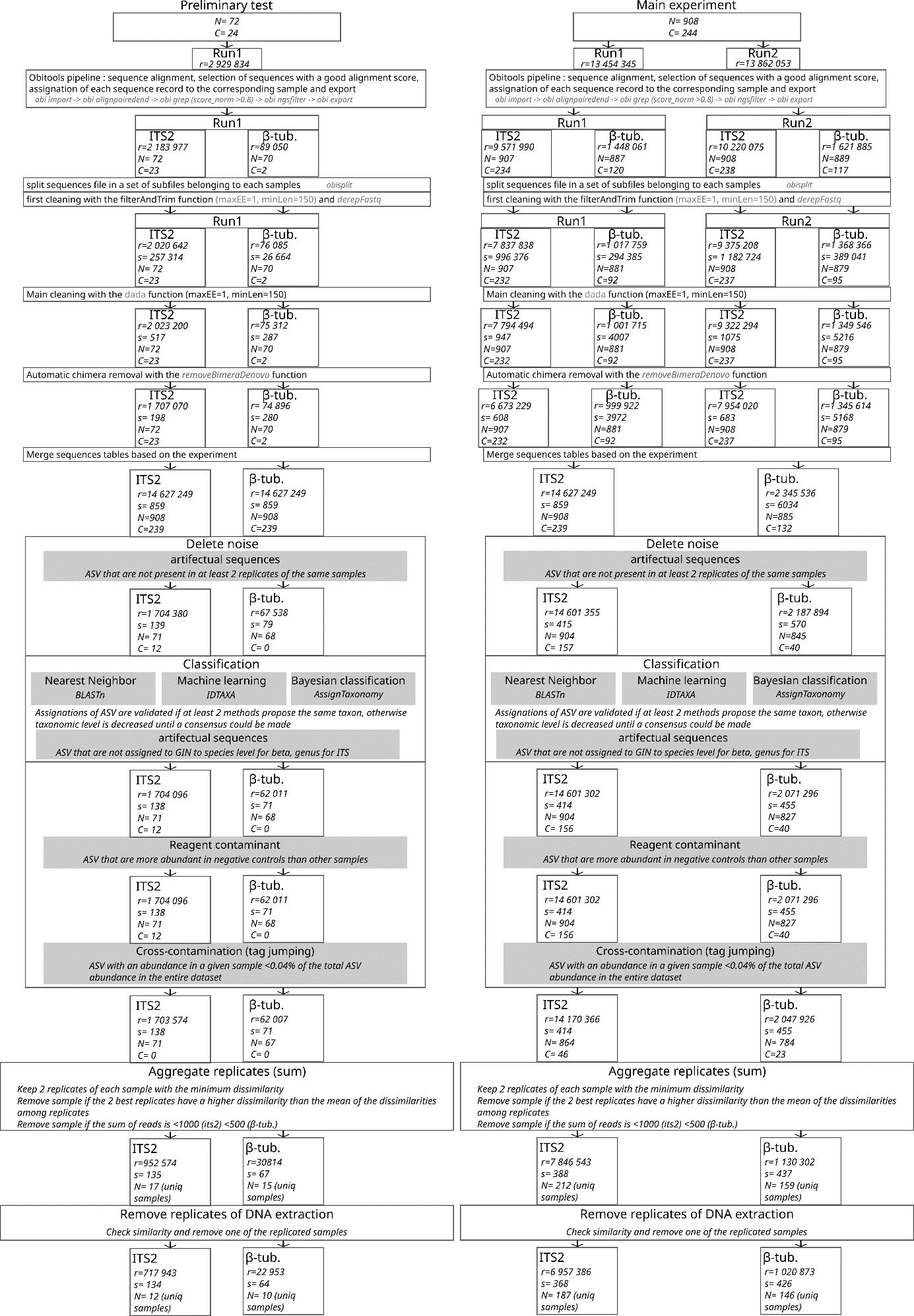

Figure S3. Workflow of the sequencing data analysis. R: reads, s: sequences, N: biological samples, C: control samples.

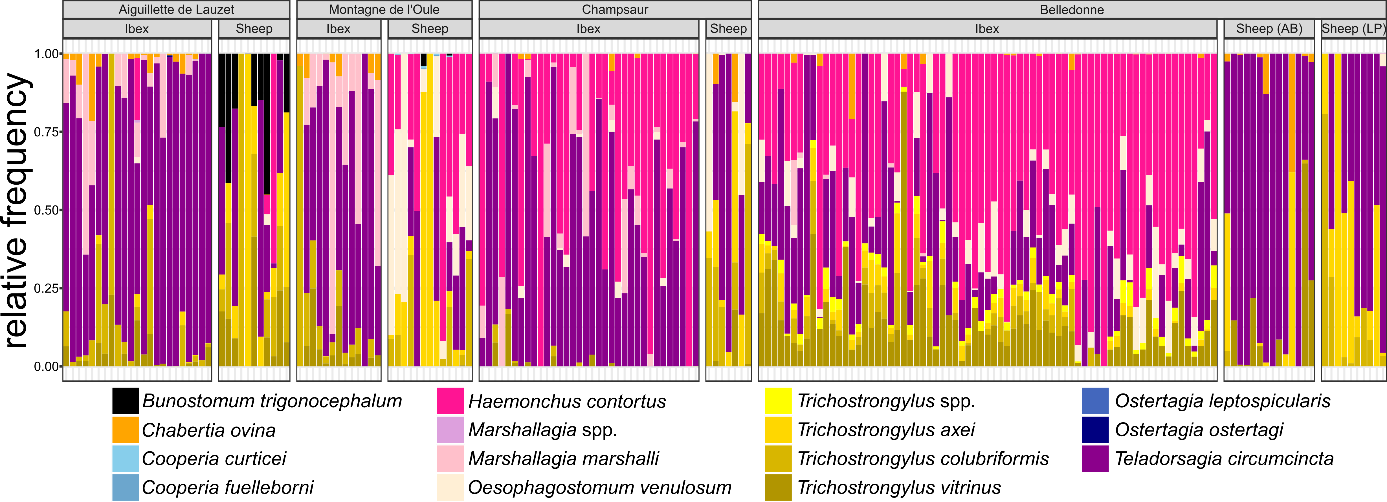

Figure S4. Read relative frequencies of gastrointestinal nematodes at taxa level. Each stacked bar chart represents the species composition of ibex and sheep samples within which each taxon is defined by one color. The data are split based on site location and species.

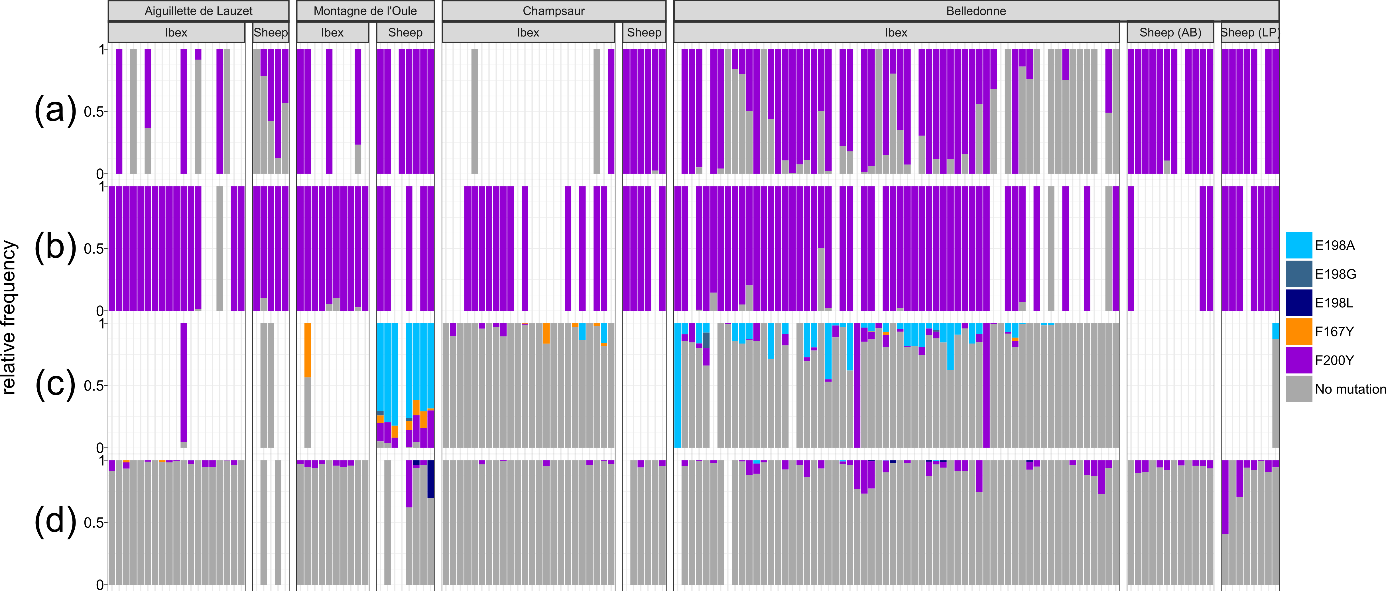

Figure S5. Read relative frequencies of resistant strains of gastrointestinal nematodes. Each stacked bar chart represents the relative proportion of isotype-1 β-tubulin resistance allele frequencies of ibex and sheep samples for four different trichostrongylid nematode species. (a) *T. axei* ; (b) *T. colubriformis* ; (c) *H. contortus* ; (d) *T. circumcincta*.

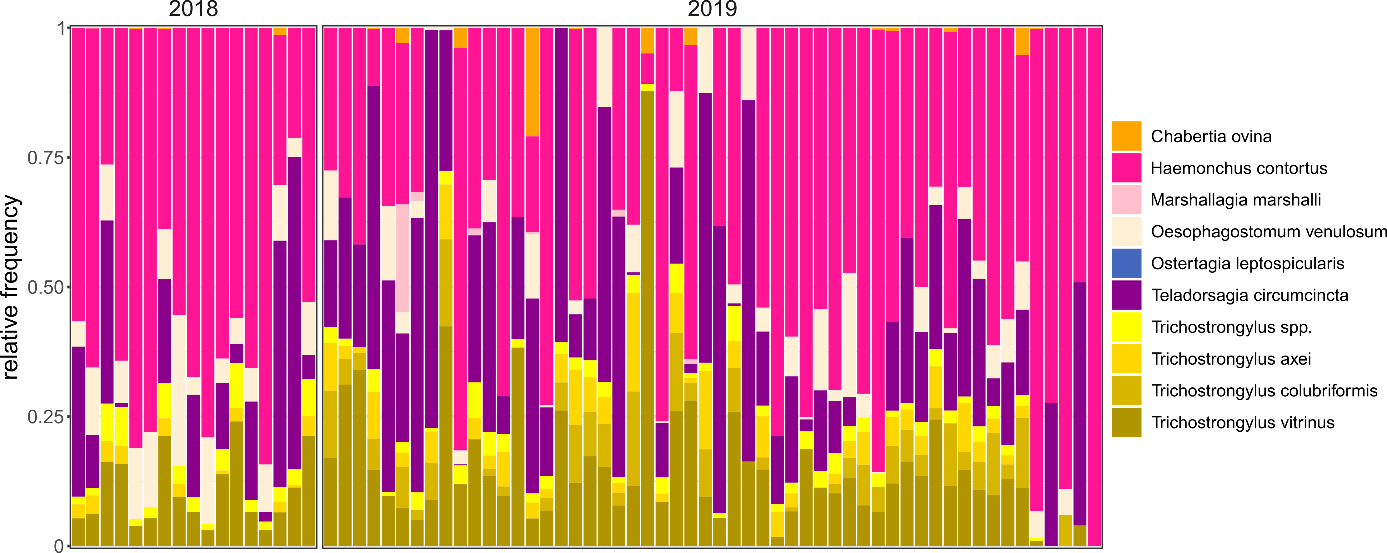
**Figure S6**. Nemabiome of ibex in Belledonne. Each stacked bar chart represents the species composition of ibex samples within which each taxon is defined by one color. The data are split based on year of sampling.

Table S1. Set of generalized linear models explaining the alpha diversity sorted by AICc value. The best model is highlighted in bold (i.e., the most parsimonious model among those with ΔAICc≤2).

| Generalized linear models | df | AICc | ΔAICc | weight |
| --- | --- | --- | --- | --- |
| $\boldsymbol{\alpha\sim species}$ | **3** | **239.7** | **0** | **0.62** |
| $\boldsymbol{\alpha\sim site+species}$ | 6 | 240.9 | 1.18 | 0.34 |
| $\boldsymbol{\alpha\sim site+species+site\times species}$ | 9 | 245.4 | 5.70 | 0.04 |
| $\boldsymbol{\alpha\sim site}$ | 5 | 267.4. | 27.74 | 0 |
| $\boldsymbol{\alpha\sim1}$ | 2 | 270.3 | 30.61 | 0 |

Table S2. Set of perMANOVA models explaining the beta diversity sorted by AICc value. The best model is highlighted in bold (i.e., the most parsimonious model among those with ΔAICc≤2).

| perMANOVA models | k | AICc | ΔAICc | weight |
| --- | --- | --- | --- | --- |
| $\boldsymbol{\beta\sim site+species+site\times species}$ | **8** | **-463.88** | **0** | **1** |
| $\boldsymbol{\beta\sim site+species}$ | 5 | -403.17 | 60.71 | 0 |
| $\boldsymbol{\beta\sim site}$ | 4 | -385.78 | 78.10 | 0 |
| $\boldsymbol{\beta\sim species}$ | 2 | -380.87 | 83.01 | 0 |
| $\boldsymbol{\beta\sim1}$ | 1 | 642.54 | 1106.42 | 0 |

Table S3. Set of generalized linear models explaining the alpha diversity sorted by AICc value. The best model is highlighted in bold (i.e., the most parsimonious model among those with ΔAICc≤2).

| Generalized linear models | df | AICc | ΔAICc | weight |
| --- | --- | --- | --- | --- |
| $\boldsymbol{\alpha}\boldsymbol{\sim}\boldsymbol{site}\boldsymbol{+}\boldsymbol{classes}\boldsymbol{+}\boldsymbol{site}\boldsymbol{\times classes}$ | **7** | **40.24** | **0** | **0.72** |
| $\boldsymbol{\alpha\sim site+classes}$ | 5 | 42.41 | 2.16 | 0.25 |
| $\boldsymbol{\alpha\sim classes}$ | 3 | 48.04 | 7.80 | 0.01 |
| $\boldsymbol{\alpha\sim1}$ | 2 | 49.16 | 8.92 | 0.01 |
| $\boldsymbol{\alpha\sim site}$ | 4 | 49.39 | 9.15 | 0.01 |

Table S4. Set of perMANOVA models explaining the beta diversity sorted by AICc value. The best model is highlighted in bold (i.e., the most parsimonious model among those with ΔAICc≤2).

| perMANOVA models | k | AICc | ΔAICc | weight |
| --- | --- | --- | --- | --- |
| $\boldsymbol{\beta\sim site}$ | **3** | **-187.61** | **0** | **0.45** |
| $\boldsymbol{\beta\sim site+classes}$ | 4 | -187.39 | 0.22 | 0.40 |
| $\boldsymbol{\beta}\boldsymbol{\sim}\boldsymbol{site}\boldsymbol{+classes+}\boldsymbol{site}\boldsymbol{\times classes}$ | 6 | -185.47 | 2.14 | 0.15 |
| $\boldsymbol{\beta\sim classes}$ | 2 | -168.26 | 19.35 | 0 |
| $\boldsymbol{\beta\sim1}$ | 1 | 79.39 | 267.00 | 0 |

Table S5. Set of generalized linear models explaining model explaining the resistant reads relative abundance (RRA) in ibex and sheep sorted by AICc value. The best model is highlighted in bold (i.e., the most parsimonious model among those with ΔAICc≤2). GIN : gastrointestinal nematode.

| Generalized linear models | df | AICc | ΔAICc | weight |
| --- | --- | --- | --- | --- |
| $\boldsymbol{\alpha\sim host species+GIN species+site+host species\times site}$ | **11** | **215.8** | **0** | **0.68** |
| $\boldsymbol{\alpha\sim host species+GIN species+site+host species\times GIN species+ host species\times site}$ | 14 | 218.3 | 2.56 | 0.19 |
| $\boldsymbol{\alpha\sim host species+GIN species+site}$ | 8 | 219.4 | 3.58 | 0.11 |
| $\boldsymbol{\alpha\sim host species+GIN species+site+host species\times GIN species}$ | 11 | 225.5 | 9.71 | 0.01 |
| $\boldsymbol{\alpha\sim host species+GIN species+site+host species\times site+site\times GIN species+host species\times GIN species}$ | 23 | 226.2 | 10.42 | 0 |
| $\boldsymbol{\alpha\sim host species+GIN species+site+host species\times site+site\times GIN species}$ | 20 | 226.2 | 10.43 | 0 |
| $\boldsymbol{\alpha\sim host species+GIN species}$ | 5 | 227.0 | 11.18 | 0 |
| $\boldsymbol{\alpha\sim host species+GIN species+site+site\times GIN species}$ | 17 | 230.4 | 14.57 | 0 |
| $\boldsymbol{\alpha\sim host species+GIN species+host species\times GIN species}$ | 8 | 230.7 | 14.95 | 0 |
| $\boldsymbol{\alpha\sim host species+GIN species+site+site\times GIN species+host species\times GIN species}$ | 20 | 236.3 | 20.54 | 0 |
| $\boldsymbol{\alpha\sim GIN species+site}$ | 7 | 237.0 | 21.25 | 0 |
| $\boldsymbol{\alpha\sim GIN species+site+site\times GIN species}$ | 16 | 241.5 | 25.75 | 0 |
| $\boldsymbol{\alpha\sim host species+GIN species+site+host species\times site+site\times GIN species+host species\times GIN species+host species\times GIN species\times site}$ | 31 | 242.9 | 27.08 | 0 |
| $\boldsymbol{\alpha\sim GIN species}$ | 4 | 259.4 | 43.58 | 0 |
| $\boldsymbol{\alpha\sim host species+site}$ | 5 | 597.5 | 381.74 | 0 |
| $\boldsymbol{\alpha\sim host species+site+host species\times site}$ | 8 | 598.3 | 382.49 | 0 |
| $\boldsymbol{\alpha\sim host species}$ | 2 | 603.1 | 387.35 | 0 |
| $\boldsymbol{\alpha\sim site}$ | 4 | 614.5 | 398.70 | 0 |
| $\boldsymbol{\alpha\sim1}$ | 1 | 626.3 | 410.51 | 0 |
